# TFAP2C is a key regulator of intrauterine trophoblast cell invasion and deep hemochorial placentation

**DOI:** 10.1101/2024.10.31.621324

**Authors:** Esteban M. Dominguez, Ayelen Moreno-Irusta, Regan L. Scott, Khursheed Iqbal, Michael J. Soares

## Abstract

Transcription factor AP-2 gamma (**TFAP2C**) has been identified as a key regulator of the trophoblast cell lineage and hemochorial placentation. The rat possesses deep placentation characterized by extensive intrauterine trophoblast cell invasion, which resembles human placentation. *Tfap2c is* expressed in multiple trophoblast cell lineages, including invasive trophoblast cells situated within the uterine-placental interface of the rat placentation site. Global genome-editing was used to explore the biology of *Tfap2c* in rat placenta development. Homozygous global disruption of *Tfap2c* resulted in prenatal lethality. Heterozygous global disruption of *Tfap2c* was associated with diminished invasive trophoblast cell infiltration into the uterus. The role of TFAP2C in the invasive trophoblast cell lineage was explored using Cre-lox conditional mutagenesis. Invasive trophoblast cell-specific disruption of *Tfap2c* resulted in inhibition of intrauterine trophoblast cell invasion and intrauterine and postnatal growth restriction. The invasive trophoblast cell lineage was not impaired following conditional monoallelic disruption of *Tfap2c*. In summary, TFAP2C contributes to the progression of distinct stages of placental development. TFAP2C is a driver of early events in trophoblast cell development and reappears later in gestation as an essential regulator of the invasive trophoblast cell lineage. A subset of TFAP2C actions on trophoblast cells are dependent on gene dosage.

## Introduction

The placenta is a transient organ that ensures viviparity and optimal fetal development (1–3). The human and rat possess a hemochorial placenta (4, 5), which is composed of specialized cellular constituents termed trophoblast cells (6). Among the trophoblast cell lineages contributing to the placentation site are invasive trophoblast cells, which in the human are referred to as extravillous trophoblast cells (7). Human and rat placentation sites are characterized by deep intrauterine trophoblast cell invasion (8–10). Invasive trophoblast cells facilitate the erosion of maternal uterine spiral arteries permitting a direct flow of maternal nutrients to trophoblast cells (8, 11–15). Uterine vasculature remodeling is a critical process ensuring a successful pregnancy outcome (13, 16, 17). There is a paucity of knowledge regarding regulatory mechanisms controlling development of the invasive trophoblast cell lineage and trophoblast cell-guided uterine spiral remodeling.

Members of the transcription factor AP-2 (**TFAP2**) family are key regulators of embryonic and extraembryonic development (18, 19). TFAP2C distinguishes itself as a regulator of the trophoblast cell lineage and hemochorial placentation (20–22). In the mouse, global disruption of the *Tfap2c* locus results in abnormalities within the trophoblast cell lineage and prenatal lethality prior to placentation (23, 24). Although, TFAP2C is an established regulator of the earliest stages of trophoblast cell lineage development (25–28), placental expression profiles and target gene occupancy implicates TFAP2C in the regulation of multiple processes impacting mammalian placentation and placental function (29–40). Included in this broad spectrum of TFAP2C actions on placentation is its potential involvement in invasive trophoblast cell biology (38–41).

In this report, we present the results of in vivo experimentation directed toward elucidating the physiological role of TFAP2C in the developing hemochorial placenta with a focus on trophoblast cell-guided events within the uterine-placental interface. We show that TFAP2C possesses essential roles during distinct stages of placental development. TFAP2C is an essential regulator of early placental morphogenetic events and reappears in a critical capacity for invasive trophoblast cell lineage development. Finally, elements of TFAP2C actions on placentation are dose dependent.

## Results

### TFAP2C expression within the placentation site

The rat hemochorial placenta is comprised of multiple lineages of trophoblast cells arranged into distinct compartments, including the labyrinth zone, junctional zone, and uterine-placental interface (42, 43) (Figure 1). The labyrinth zone is the site of maternal-fetal nutrient/waste transfer (44), whereas the junctional zone is composed of endocrine cells targeting the maternal environment and the site where invasive trophoblast cells arise (42, 43, 45). Cell composition of the uterine-placental interface is dynamic, including an increase in the infiltration of invasive trophoblast cells as gestation proceeds (9, 38). Intrauterine invasive trophoblast cells invade into the stroma located between uterine blood vessels (interstitial) and within uterine blood vessels (endovascular). *Tfap2c* transcripts were detected within the uterine-placental interface and increased during pregnancy (Supplemental Figure 1A). In situ hybridization was used to determine the distribution of *Tfap2c* transcripts within the hemochorial placenta. At gestation day (**gd**) 9.5, *Tfap2c* was prominently expressed in the ectoplacental cone (Figure 1A). *Tfap2c* transcripts were also localized to the anti-mesometrial uterine deciduum (Figure 1A). As gestation advanced (gd 11.5), *Tfap2c* expression was observed within trophoblast cells of the developing placenta (Figure 1B). On gd 18.5, *Tfap2c* was abundantly expressed within the junctional zone and in invasive trophoblast cells of the uterine-placental interface (Figure 1, C-E). The presence of *Tfap2c* transcripts in the trophoblast cell lineage was supported by its colocalization with keratin 8 (***Krt8***), a pan-trophoblast cell marker (Figure 1, A-C). Expression of *Tfap2c* transcripts in invasive trophoblast cells was further demonstrated by co-localization with prolactin family 7, subfamily b, member 1 (*Prl7b1*) transcripts (Figure 1, D and E), an invasive trophoblast cell-specific transcript, and is consistent with single cell RNA-sequencing of the of the uterine-placental interface (Supplemental Figure 1B) (38). TFAP2C protein was distributed in invasive trophoblast cells of the uterine-placental interface and throughout the junctional zone, replicating the distribution of *Tfap2c* transcripts at gd 18.5 (Supplemental Figure 2). Within the invasive trophoblast cell lineage, TFAP2C transcript and protein were localized to both endovascular and interstitial trophoblast cells (Figure 1E, Supplemental Figure 3). *Tfap2c* transcripts were localized to most trophoblast cell populations within the junctional zone, including a smaller subset of invasive trophoblast progenitor cell populations (*Prl7b1* positive; Figure 1F).

**Figure 1.**
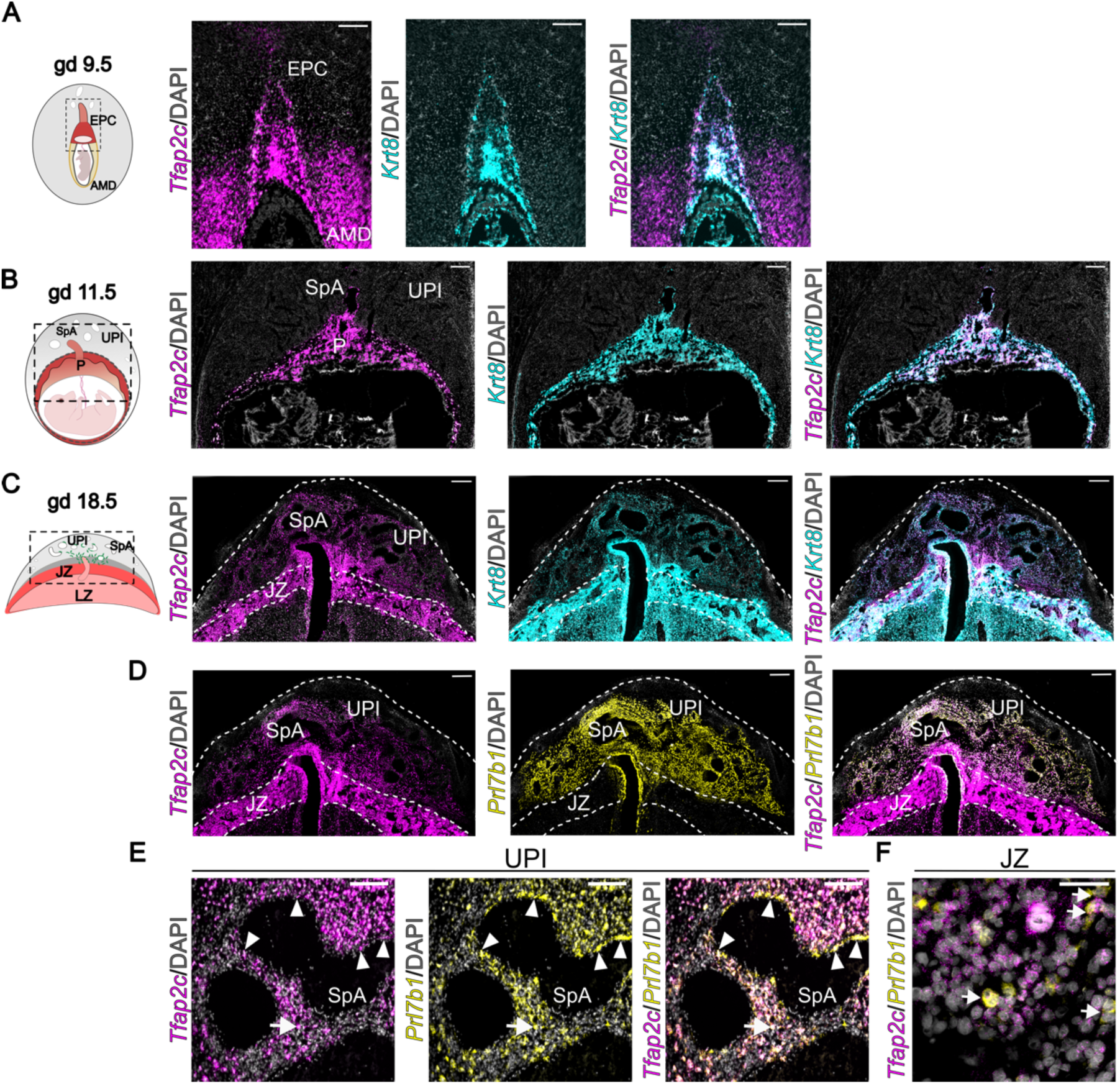
*Tfap2c* transcript distribution during placenta development. (**A-C**) *Tfap2c* transcripts (magenta) were localized to the ectoplacental cone at gestation day (gd) 9.5 (**A**) and placentation sites at 11.5 (**B**), and 18.5 (**C**). *Tfap2c* transcripts were co-localized with *Krt8* transcripts (cyan). Schematic diagrams representing gd 9.5, 11.5, and 18.5 placentation sites are initially presented for each panel. (**D**) Co-localization of *Tfap2c* (magenta) and *Prl7b1* (yellow) transcripts within the gd 18.5 placentation site. *Prl7b1* is specifically expressed in the invasive trophoblast cell lineage. Scale bar: 500 μm. (**E**) High magnification images showing *Tfap2c* and *Prl7b1* transcript localization in endovascular (arrowhead) and interstitial (arrow) invasive trophoblast cells within the uterine-placental interface. Scale bar: 100 μm. (**F**) Co-localization of *Tfap2c* and *Prl7b1* transcripts (arrows) within the junctional zone of the gd 15.5 placenta. Scale bar: 50 μm. Co-localization is shown as whilte. Abbreviations: EPC, ectoplacental cone; AMD, anti-mesometrial decidua; SpA, spiral artery; UPI, uterine-placental interface; P, placenta; JZ, junctional zone; LZ, labyrinth zone.

### Global deletion of *Tfap2c* leads to prenatal lethality

Next, we investigated the in vivo role of *Tfap2c* in rat placenta development using CRISPR/Cas9 genome editing. DNA-binding and dimerization domains encoded by Exon 4 of the *Tfap2c* gene were targeted (Supplemental Figure 4A). A germ line mutant rat was generated possessing a 308-bp deletion within Exon 4 (Supplemental Figure 4B). The deletion resulted in a frameshift and a premature stop codon (Supplemental Figure 4C). Polymerase chain reaction (**PCR**) products corresponding to wild type and mutant alleles were identified by genotyping (828 bp versus 520 bp, respectively, Supplemental Figure 4D). Null *Tfap2c* conceptuses (-/-) were obtained by mating heterozygous (+/-) *Tfap2*c females and males. Breeding results collected throughout pregnancy and postnatally, including Mendelian ratios, are presented in Supplemental Table 1. Null *Tfap2c* conceptuses were viable until gd 8.5, which temporally coincides with the onset of placental morphogenesis. This observation agrees with the timing of embryonic demise observed following global *Tfap2c* mouse gene disruption (23, 24). Interestingly, we observed an unexpected reduction in the number of surviving heterozygous pups generated from *Tfap2c^+/-^*x *Tfap2c^+/-^*, which prompted an investigation of placental development in conceptuses possessing a single *Tfap2c* mutant allele.

### Decreased *Tfap2*c gene dosage leads to a failure in intrauterine trophoblast cell invasion

To explore possible *Tfap2c* gene dosage effects, *Tfap2c^+/+^* females and *Tfap2c^+/-^* males were mated, and pregnancies assessed at gd 18.5. *Tfap2c^+/+^* and *Tfap2c^+/-^* placentation sites exhibited a similar organizational structure, including a labyrinth zone, junctional zone, and a uterine-placental interface (Figure 2A). We next assessed infiltration of invasive trophoblast cells into the uterine-placental interfaces of *Tfap2c^+/+^* versus *Tfap2c^+/-^* placentation sites by monitoring the distribution of cytokeratin protein expression (9, 38) (Figure 2B) and *Prl7b1* mRNA (46) (Figure 2C), both representing indices of invasive trophoblast cells. The uterine-placental interface of *Tfap2c^+/-^*placentation sites were depleted of invasive trophoblast cells (Figure 2, B and C). Both interstitial and endovascular invasive trophoblast cells were diminished in the uterine-placental interface of *Tfap2c^+/-^* placentation sites. This result was supported by quantification of the depth of invasion of cytokeratin positive cells into the uterine parenchyma (Figure 2D). The depletion of invasive trophoblast cells in the uterine-placental interface of *Tfap2c^+/-^* placentation sites was further supported by diminished expression of several invasive trophoblast cell associated transcripts (*Krt7, Krt8, Krt18, Prl7b1, Prl5a1, Cdkn1c, Cited2, Plac1, Peg3, Tfpi;* (38) (Figure 2E). Similar findings were recorded when placentation sites were interrogated from pregnancies generated from *Tfap2c^+/-^* females mated to *Tfap2c^+/+^* males (Supplemental Figure 5, A-D).

**Figure 2.**
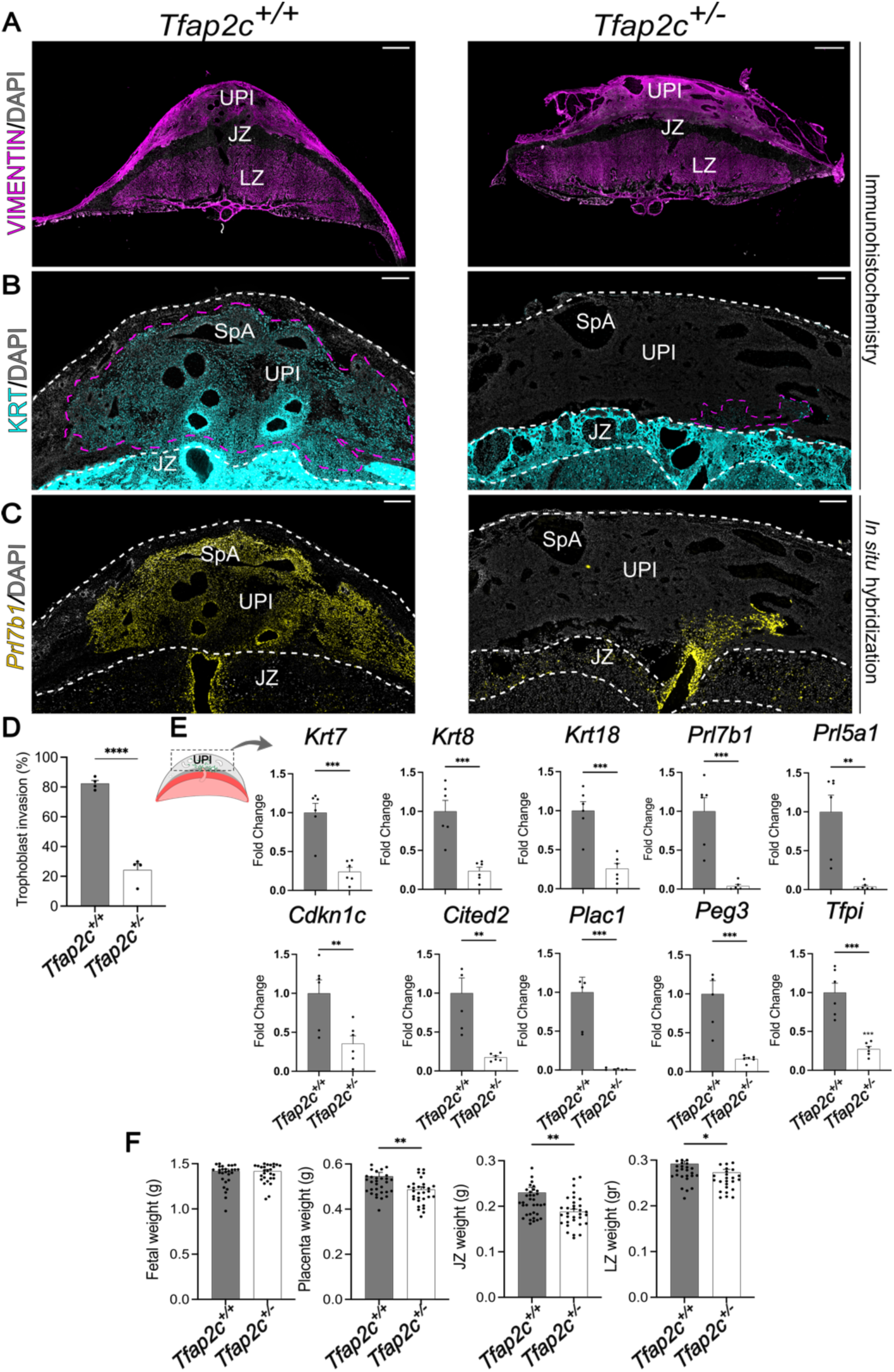
*Tfap2c* gene dosage effects on placental development. Placentation sites were generated from *Tfap2c^+/-^* male and *Tfap2c^+/+^* female breeding. (**A**) Identification of placentation site compartments *Tfap2c^+/+^* and *Tfap2c^+/-^* using vimentin immunostaining (magenta). (**B**) Immunohistochemical localization of cytokeratin protein in gestation day (gd) 18.5 *Tfap2c^+/+^* and *Tfap2c^+/-^* placentation sites (cyan). (**C**) Distribution of *Prl7b1* transcripts in gd 18.5 *Tfap2c^+/+^* and *Tfap2c^+/-^* placentation sites (yellow). (**D**) Quantification of trophoblast cell invasion area (magenta dotted line) determined by cytokeratin immunostaining within the uterine-placental interface. Data are expressed as mean ± standard error of the mean (SEM). Each data point represents different uterine-placental interface tissues (n=4) obtained from four pregnancies. (**E**) RT-qPCR of invasive trophoblast cell-specific transcripts in gd 18.5 *Tfap2c^+/+^* and *Tfap2c^+/-^* uterine-placental interface tissues. Data are expressed as mean ± SEM. Each data point represents a biological replicate obtained from six different pregnancies (n=6). (**F**) Fetal, placenta, junctional zone and labyrinth zone weights for gd 18.5 *Tfap2c^+/+^* and *Tfap2c^+/-^* conceptuses. Data are expressed as mean ± SEM. Each data point represents a biological replicate obtained from six different pregnancies (*Tfap2+^+/+^*, n=38; *Tfap2c ^+/-^*, n=32). Unpaired *t*-test: * p<0.05, **p<0.01, ***p<0.001, ****p<0.0001. Abbreviations: UPI, uterine-placental interface; JZ, junctional zone; LZ, labyrinth zone; SpA, spiral artery. Scale bars: 500 μm.

However, breeding combinations did differentially affect postnatal heterozygote offspring viability and heterozygote fetal and placental weights. Postnatal heterozygote viability was prominently decreased in female *Tfap2c^+/-^* x male *Tfap2c^+/-^* but not in female *Tfap2c^+/+^* x male *Tfap2c^+/-^* or female *Tfap2c^+/-^* x male *Tfap2c^+/+^*breeding combinations (Supplemental Table 1-3). Genotypic differences in fetal, placenta, junctional zone, and labyrinth zone weights at gd 18.5 were observed among breeding combinations. Mating *Tfap2c^+/+^* females with *Tfap2c^+/-^* males was associated with significant decreases in heterozygote placenta, junctional zone, and labyrinth zone weights (Figure 2F), whereas only heterozygote fetal weights were significantly decreased by mating *Tfap2c^+/-^*females with *Tfap2c^+/+^* males (Supplemental Figure 5E). Some of these differences might be associated with *Tfap2c* gene dosage effects on the maternal environment.

### Invasive trophoblast cell conditional *Tfap2c* mutagenesis

The impact of TFAP2C dosage on invasive trophoblast cell development prompted a direct analysis of the importance of TFAP2C on the invasive trophoblast cell lineage. We generated an invasive trophoblast cell-specific *Tfap2c* gene disruption using the Cre/loxP system. CRISPR/Cas9 genome editing was used to insert loxP sites flanking Exon 4 of the *Tfap2c* gene (Figure 3A). *Tfap2c* floxed (***Tfap2c^f/f^***) rats were mated with *Prl7b1-Cre* rats (47). Cre recombinase is specifically targeted to invasive trophoblast cells in *Prl7b1-Cre* rats (47). TFAP2C protein was present in lysates of the *Tfap2c^f/f^* uterine-placental interface but not in lysates of the *Tfap2c^d/d^* uterine-placental interface (Figure 3B). Immunostaining supported depletion of TFAP2C from the gd 18.5 uterine-placental interface but not from the JZ of *Tfap2c^d/d^* placentation sites (Figure 3C). Our ability to modulate TFAP2C in the invasive trophoblast cell lineage led us to evaluate a potential role for TFAP2C in regulating the invasive trophoblast cell lineage.

**Figure 3.**
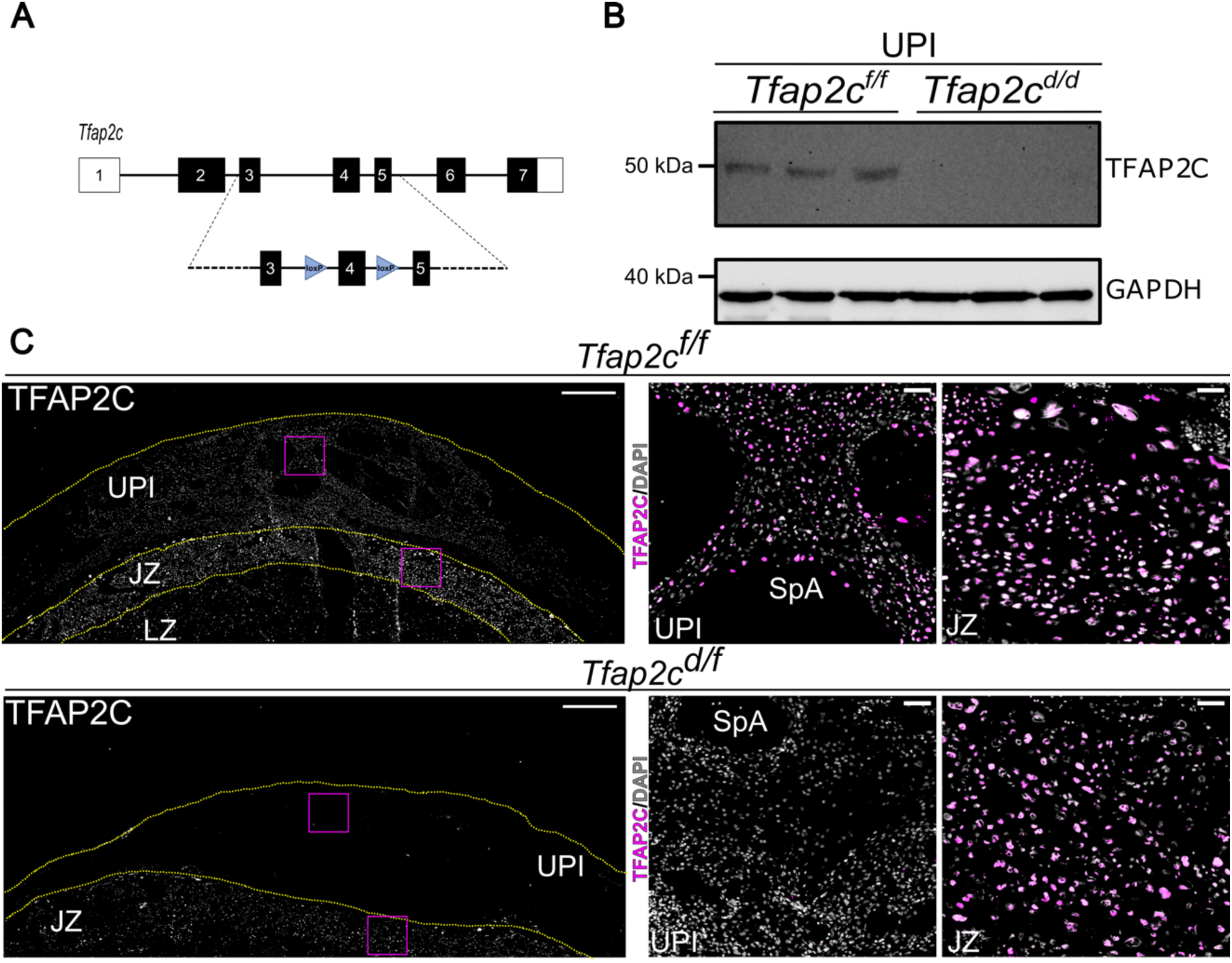
Generation of an invasive trophoblast cell-specific *Tfap2c* deletion. (**A**) Schematic of *loxP* sites inserted into introns flanking Exon 4 of the *Tfap2c* gene. (**B**) TFAP2C western blot of the uterine-placental interface of *Tfap2c^f/f^* and *Tfap2c^d/d^* gd 18.5 placentation sites. (**C**) TFAP2C immunostaining of *Tfap2c^f/f^* and *Tfap2c^d/d^* gd 18.5 placentation sites (left panels: TFAP2C immunostaining, white; right panels: TFAP2C immunostaining, magenta, DAPI, white). Scale bar: 500 μm. Inserts to the right show higher magnification images of uterine-placental interface tissues (top) and junctional zone tissue (bottom) for each placentation site. Scale bar: 100 μm. Abbreviations: UPI, uterine-placental interface; JZ, junctional zone; LZ, labyrinth zone; SpA, spiral artery.

### TFAP2C-dependent intrauterine trophoblast cell invasion

Intrauterine trophoblast cell invasion was assessed in *Tfap2c^f/f^*and *Tfap2c^d/d^* placentation sites. At gd 18.5 invasive trophoblast cells were present throughout the *Tfap2c^f/f^* uterine-placental interface but were not detected in the *Tfap2^d/d^* uterine-placental interface (Figure 4, A-C). Interstitial and endovascular invasive trophoblast cells were absent from the uterine-placental interface of *Tfap2c^d/d^* placentation sites. The depletion of invasive trophoblast cells in the uterine-placental interface of *Tfap2c^d/d^* placentation sites was further supported by diminished expression of several invasive trophoblast cell-associated transcripts (*Krt7, Krt8, Krt18, Prl7b1, Prl5a1, Cdkn1c, Cited2, Plac1, Peg3, Tfpi*; Figure 4D). In contrast to the global disruption of *Tfap2c*, migration of invasive trophoblast cells into the uterus was not impaired following conditional monoallelic disruption of *Tfap2c* (Supplemental Figure 6). Single cell RNA-sequencing and single nucleus assay for transposase-accessible chromatin-sequencing datasets (38, 41) reveal a network of genes potentially regulated by TFAP2C in rat invasive trophoblast cells that, include candidate contributors to remodeling extracellular matrix, cytoskeleton restructuring, regulation of cell movement, and transcriptional regulators that target genes encoding other proteins affecting the invasive trophoblast phenotype (Table 1; Supplemental Dataset 1). Thus, TFAP2C has a fundamental role in regulating the invasive trophoblast cell lineage, especially movement of invasive trophoblast cells into the uterus, and a list of downstream effectors are available to investigate mechanisms underlying TFAP2C actions.

**Figure 4.**
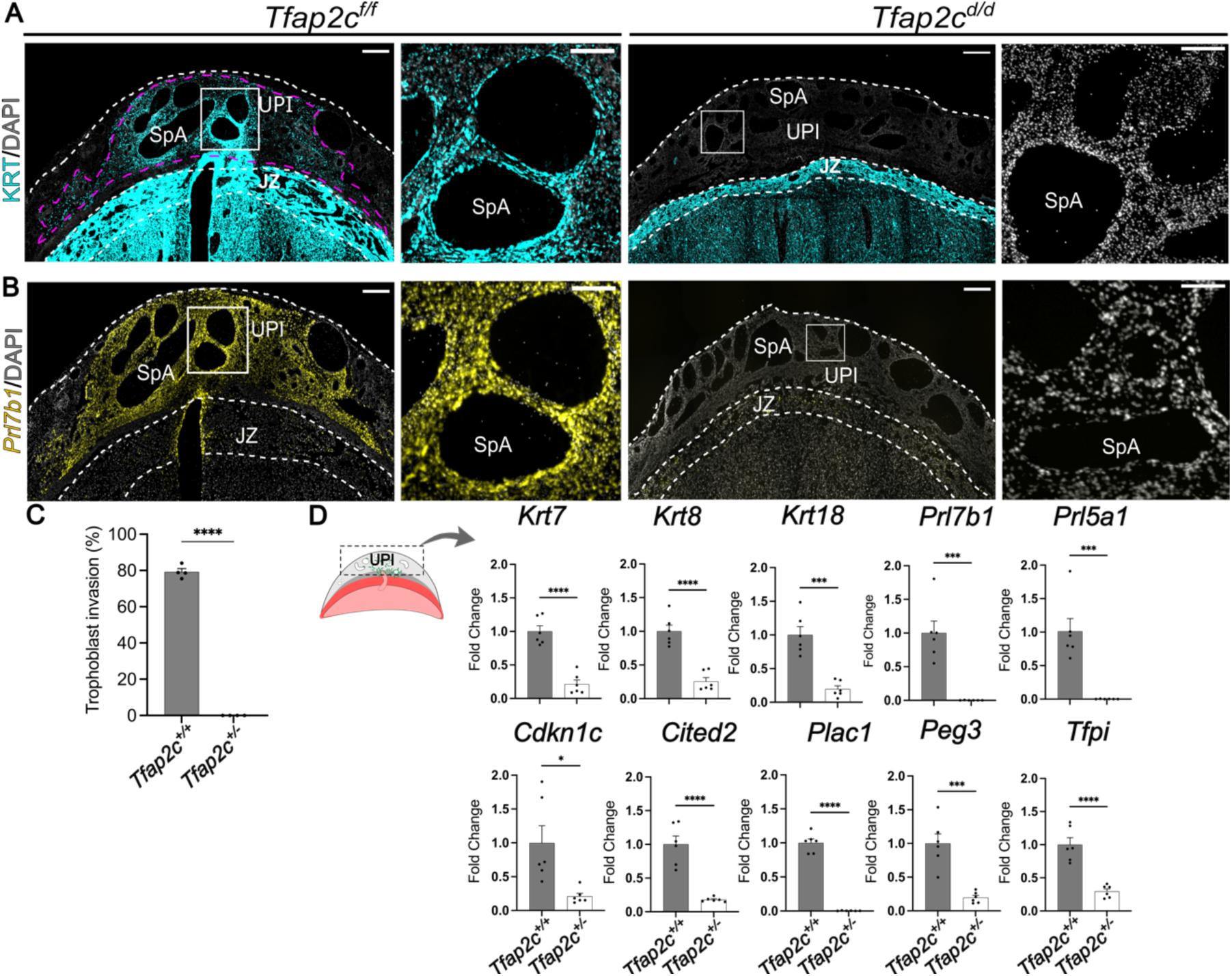
Characterization of invasive trophoblast cell-specific *Tfap2c* deletion. (**A**) Immunohistochemical localization of cytokeratin (magenta) in gd 18.5 *Tfap2c^f/f^* and *Tfap2c ^d/d^* placentation sites. (**B**) Distribution of *Prl7b1* transcripts (yellow) in gd 18.5 *Tfap2c ^f/f^* and *Tfap2c ^d/d^* placentation sites. Scale bar: 500 μm. Inserts to the right show higher magnification images of the boxed areas within the uterine-placental interface tissues for each placentation site. Scale bar: 100 μm. (**C**) Quantification of trophoblast cell invasion area (green dotted line) determined by cytokeratin immunostaining within the uterine-placental interface. Data are expressed as mean ± standard error of the mean (SEM). Each data point represents different uterine-placental interface tissues obtained from four pregnancies (n=4). (**D**) RT-qPCR of invasive trophoblast cell-specific transcripts in gd 18.5 *Tfap2c ^f/f^* and *Tfap2c ^d/d^* uterine-placental interface tissues. The left panel corresponds to a schematic of the regions of the placentation site used for analysis. Data are expressed as mean ± SEM. Each data point represents a biological replicate from six different pregnancies (n=6). Unpaired *t*-test: *p< 0.05, ***p<0.001, ****p<0.0001.). Abbreviations: UPI, uterine-placental interface; JZ, junctional zone; LZ, labyrinth zone; SpA, spiral artery.

**Table 1.** Subset of genes expressed in invasive trophoblast cells possessing TFAP2C DNA binding motifs.

| **Extracellular matrix organization** | **Cell movement - migration** | **Cytoskeletal organization** | **Transcriptional regulation** |
| --- | --- | --- | --- |
| Adamts3, Adgrg6<br>Col4a1<br>Fbln1<br>Galnt6<br>Lama4<br>Lamb1<br>Nid1<br>Tp53i11 | Amotl2, Arhagp8,<br>Cald1, Dab2ip<br>Efnb1, Fbln1<br>Mta3, Ncam1<br>Nuak1, Parva<br>Pcdh12, Plet1<br>Sema6d, Septin4<br>Svil, Tbc1d9 | Ankrd17, Apbb1ip<br>Cald1, Cfl2<br>Daam1<br>Dsp<br>Klhl2<br>Krt7<br>Krt8<br>Svil | Barx2, Bend5<br>Cdx2, Cited2<br>Ets2, Hdac4<br>Hic2, Jdp2<br>Ncor2, Nr2f6<br>Nrip1, Peg3<br>Pparg, Tfap2c<br>Zfp706 |

### Invasive trophoblast cells and natural killer (NK) cells exhibit a reciprocal distribution within the uterine-placental interface

At midgestation, NK cells are an abundant constituent of the uterine-placental interface (48). As gestation progresses, NK cells become depleted as trophoblast cells invade the uterine-placental interface (48). Interestingly, we observed a retention of NK cells in *Tfap2c^d/d^* uterine-placental interfaces depleted of invasive trophoblast cells (Figure 5A). This observation was further supported by increased expression of NK cell-associated transcripts (*Prf1, Klrb1c, Klrb1a, Klrb1, Ncr1*; Figure 5B) within *Tfap2c^d/d^* uterine-placental interfaces. These findings reinforce the observation that fundamental changes occur in the *Tfap2c^d/d^* uterine-placental interface and imply that invasive trophoblast cells affect cellular dynamics within the uterine-placental interface.

**Figure 5.**
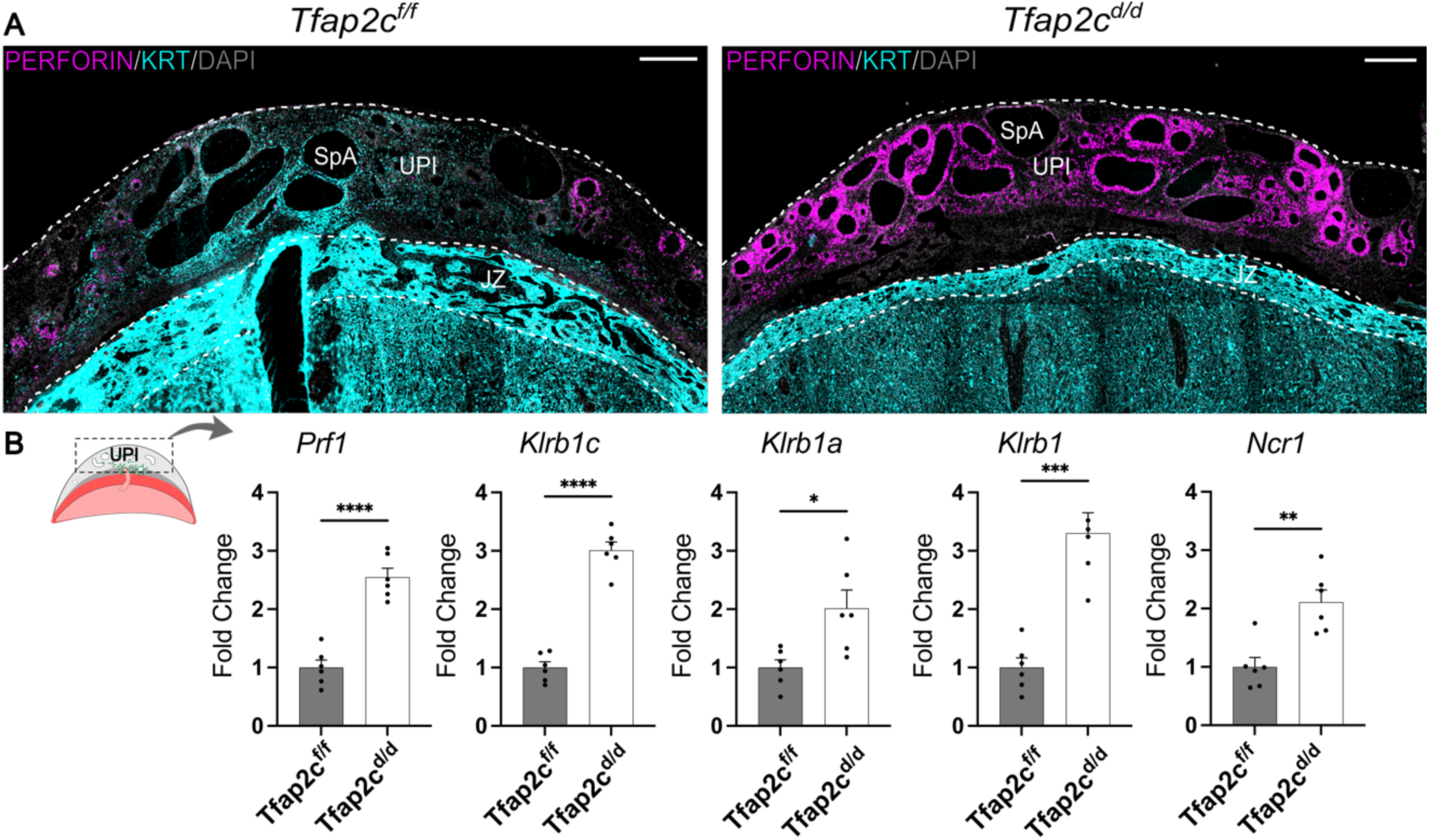
*Tfap2c* conditional deletion in trophoblast invasive cells results in the retention of uterine natural killer (NK) cells within the uterine-placental interface. (**A**) Immunohistochemical localization of cytokeratin (cyan) and perforin (magenta) in gd 18.5 *Tfap2c^f/f^* and *Tfap2c ^d/d^* placentation sites. (**B**) RT-qPCR of NK cell-specific transcripts in gd 18.5 *Tfap2c ^f/f^* and *Tfap2c ^d/d^* uterine-placental interface tissues. The left panel corresponds to a schematic of the regions of the placentation site used for analysis. Data are expressed as mean ± standard error of the mean. Each data point represents a biological replicate from six different pregnancies (n=6). Unpaired *t*-test: *p< 0.05, **p<0.01, ***p<0.001, ****p<0.0001. Abbreviations: UPI, uterine-placental interface; JZ, junctional zone; LZ, labyrinth zone; SpA, spiral artery. Scale bar: 500 μm. Specimens used in this figure are the same used in Figure 4A.

### Invasive trophoblast cells impact prenatal and postnatal growth

We next investigated whether the presence of invasive trophoblast cells within the uterine-placental interface affected prenatal or postnatal growth. Depletion of invasive trophoblast cells as observed in *Tfap2c^d/d^*placentation sites was associated with intrauterine placental growth restriction, including both junctional zone and labyrinth zone compartments, and fetal growth restriction at gd 18.5 (Figure 6A). The adverse effects of invasive trophoblast cell depletion on fetal weight extended to postnatal offspring growth. Depletion of invasive trophoblast cells was associated with decreased postnatal body weight (Figure 6B). The results are consistent with invasive trophoblast cells supporting the nutrient demands required for optimal intrauterine growth. Deficits acquired prenatally were sustained during the first three weeks of postnatal life.

**Figure 6.**
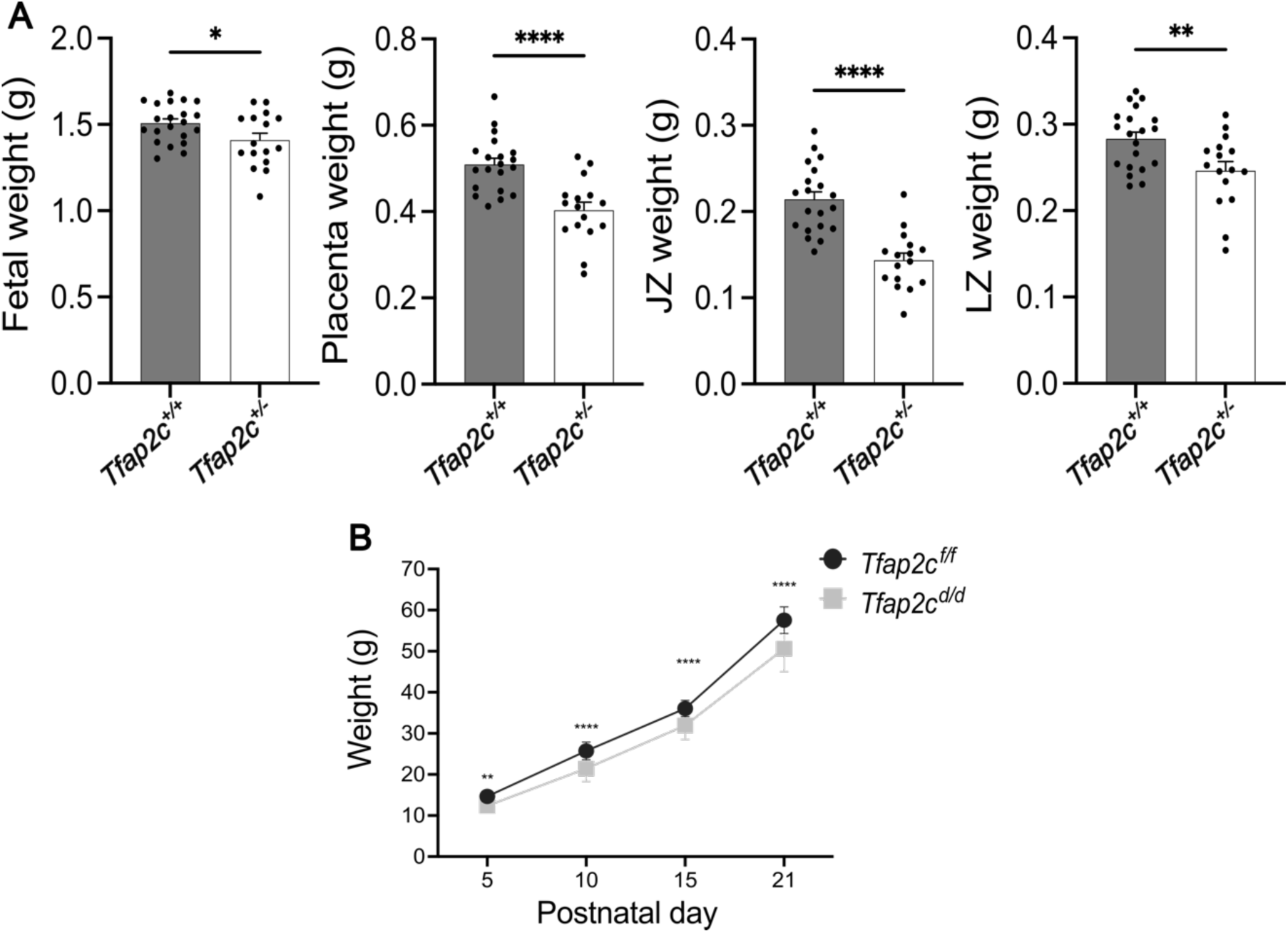
Impact of an invasive trophoblast cell-specific *Tfap2c* deletion on prenatal and postnatal development. (**A**) Fetal, placenta, junctional zone, and labyrinth zone weights from gd 18.5 *Tfap2c^f/f^* and *Tfap2c^d/d^* gd 18.5 conceptuses. Data are expressed as mean ± standard error of the mean (SEM). Each data point represents a biological replicate from six pregnancies (*Tfap2c^f/f^*, n=20; *Tfap2c ^d/d^*, n= 16). Unpaired *t*-test: *p< 0.05, **p<0.01, ***p<0.001, ****p<0.0001. (**B**) Postnatal body weight for *Tfap2c^f/f^* and *Tfap2c^d/d^* pups. Data are expressed as mean ± SEM from six pregnancies (*Tfap2c^f/f^*, n=24; *Tfap2c ^d/d^*, n= 29). One-Way ANOVA and Tukey’s Test: **p<0.01, ****p<0.0001. Abbreviations: JZ, junctional zone; LZ, labyrinth zone.

## Discussion

Placentation is a critical adaptation ensuring the success of viviparity (5, 49). Gene manipulated animal models and trophoblast stem cells have provided insights into gene regulatory networks controlling trophoblast cell lineage development and placentation (21). TFAP2C figures prominently in these networks and represents a conserved regulator of placentation (20–22). In the research reported here, we utilized genome-edited rat models to elucidate roles for TFAP2C in placentation. The rat, like the human, is a species exhibiting deep intrauterine trophoblast cell invasion (8–10). TFAP2C is expressed in multiple trophoblast cell lineages of the developing rat placenta, including invasive trophoblast cells residing in the uterine-placental interface. Consistent with earlier reports in the mouse (23, 24), global disruption of *Tfap2c* in the rat led to prenatal lethality (gd 8.5 to 9.5), prior to the initiation of placenta morphogenesis. Interestingly, decreased gene dosage of *Tfap2c* (heterozygous) was compatible with pregnancy but affected placentation, including diminished infiltration of invasive trophoblast cells into the uterine-placental interface. A role for TFAP2C in the regulation of intrauterine trophoblast cell invasion was further supported using cell lineage specific mutagenesis. *Tfap2c* was conditionally disrupted in the invasive trophoblast cell lineage, which led to a uterine-placental interface devoid of invasive trophoblast cells. Impairment of intrauterine trophoblast cell invasion had consequences, including intrauterine and postnatal growth restriction.

Involvement of TFAP2C in early events dictating trophoblast cell lineage development were reinforced from phenotyping conceptuses possessing a global disruption of the *Tfap2c* locus. TFAP2C is critical to activation of the trophectoderm transcriptome (36, 50) and development of trophectoderm (25, 26, 51, 52). Trophectoderm represents the earliest stage of trophoblast cell lineage development (53, 54). The importance of TFAP2C in early trophoblast cell lineage development is further exemplified by its utilization in the direct reprogramming of fibroblasts to the trophoblast cell lineage (27, 28).

TFAP2C actions in the rat placentation site were affected by gene dosage. Monoallelic disruption of *Tfap2c* resulted in the development of a rat uterine-placental interface possessing a diminished number of invasive trophoblast cells. Observing a phenotype in *Tfap2c* global heterozygotes was intriguing but not entirely new for *Tfap2c*. Haploinsufficiency of *Tfap2c* is linked to an increased incidence of peripartum lethality (55). Surviving *Tfap2c* heterozygotes exhibit postnatal growth restriction (24), development of germ cell tumors (56), and hippocampal dysfunction (57). Placentation is also affected by *Tfap2c* haploinsufficiency (55). The labyrinth zone compartment of the mouse placenta is disorganized in *Tfp2c* heterozygotes (55). Interestingly, the invasive trophoblast cell phenotype observed in global heterozygotes reported here was not a feature of conceptuses with a monoallelic disruption of *Tfap2c* restricted to the invasive trophoblast cell lineage. An explanation for this difference is likely connected to the wide-spread involvement of TFAP2C in multiple stages of trophoblast cell differentiation (58). This is supported by TFAP2C expression extending to many more trophoblast cells in the junctional zone than express *Prl7b1*, the gene driving Cre recombinase expression in our conditional mutagenesis rat model. Global *Tfap2c* haploinsufficiency at a key stage of trophoblast cell lineage development distinct from *Prl7b1*-expressing trophoblast cells must be relevant and responsive to *Tfap2c* gene dosage. Thus, *Tfap2c* gene dosage is utilized in a differentiation stage specific manner to guide trophoblast cell development.

TFAP2C is utilized at different stages of trophoblast cell development. TFAP2C is critical to the derivation of the trophoblast cell lineage and then repurposed for later stages of placentation (58), including the regulation of invasive trophoblast cell development. TFAP2C does not act alone. Context is critical for the actions of TFAP2C. The presence of other transcription factor partners and stoichiometry influence TFAP2C function during trophoblast cell development (21, 59). A short list of potential TFAP2C partners in invasive trophoblast cell development is available (38, 41).

Lastly, we have taken transcript and open chromatin profiles (38, 41) and created a path for elucidating gene involvement in the regulation of invasive trophoblast cell lineage development and its impact on the uterine-placental interface. We provide compelling evidence that TFAP2C is critical in establishing the invasive trophoblast cell lineage. Furthermore, we effectively demonstrate that the presence of invasive trophoblast cells in the uterus has consequences. Deficits in invasive trophoblast cell guided uterine transformation resulted in intrauterine and postnatal growth restriction. Most importantly, tools are in place to establish a gene regulatory network controlling development and function of the invasive trophoblast cell lineage.

In summary, TFAP2C contributes to the progression of distinct stages of rat placental development. TFAP2C is a driver of early events in trophoblast cell development and reappears later in gestation as an essential regulator of the invasive trophoblast cell lineage. A subset of TFAP2C actions on trophoblast cells are dependent on gene dosage.

## Methods

### Sex as a biological variable

Our study examined male and female animals. Similar findings are reported for both sexes.

### Animals and tissue collection

Holtzman Sprague–Dawley rats were acquired from Envigo. Animals were maintained in an environmentally controlled facility (14 h light and 10 h dark cycle) with food and water available ad libitum. Timed pregnancies were established using virgin female rats (8 to 10 weeks of age) mated with adult male rats (>3 months of age). Mating was confirmed by the presence of sperm in the vaginal lavage and was considered gd 0.5.

Pseudopregnant female rats were generated by mating with vasectomized male rats. Detection of a seminal plug was considered day 0.5 of pseudopregnancy. Pregnant rats were euthanized on specific days of gestation. Some placentation sites were frozen intact in dry-ice cooled heptane and stored at -80 °C for subsequent histological analyses, whereas others were dissected into uterine-placental interface, junctional zone, and labyrinth zone compartments as previously described (38, 60). Dissected placentation site compartments and fetuses were weighed, frozen in liquid nitrogen, and stored at -80°C for subsequent biochemical analyses.

### Generation of global and conditional *Tfap2c* mutant rat models

A global mutation at the rat *Tfap2c* locus was generated using CRISPR/Cas9 genome editing technology (61). RNA guides targeting Exon 4 of the *Tfap2c* gene (Supplemental Figure 4, Supplemental Table 4) were assembled with *CRISPR* RNA, trans-activating *CRISPR* RNA, and Cas9 nuclease V3 (Integrated DNA Technologies). Conditional *Tfap2c* disruption by the insertion of loxP sites targeted to 5’ and 3’ introns flanking Exon 4 (Figure 3, Supplemental Table 4, and Supplemental Table 5). Electroporation of constructs were performed into one-cell rat embryos using a NEPA21 electroporator (Nepa Gene Co Ltd). Electroporated embryos were transferred to pseudopregnant rats. Offspring were screened for mutations by polymerase chain reaction (**PCR**), and the deletion confirmed by DNA sequencing (GeneWiz). A founder mutant animal was backcrossed with wild type rats to evaluate germline transmission. Genotyping and fetal sex chromosome determination were performed as previously described (47, 61, 62). DNA was extracted from tail-tip biopsies with Red Extract-N-Amp tissue PCR kit (XNAT-1000RXN, Sigma-Aldrich) and used for genotyping. Primers used for genotyping of global and conditional mutations and sex chromosome determinations are provided (Supplemental Table 6). Body weight measurements were performed on offspring generated from specific intercrosses.

### Reverse transcription-quantitative PCR (RT-qPCR)

Total RNA was isolated from tissues with TRIzol (15596018, Thermo Fisher). Extracted RNA (1 μg) was used to synthesize complementary DNA using a High-Capacity Reverse Transcription kit (4368814, Applied Biosystems) and then diluted 1:10 in water. qPCR was performed using PowerSYBR Green PCR Master Mix (4367659, Thermo Fisher) and transcript-specific primer sets (250 nM). Primer sequences are provided (Supplemental Table 7). QuantStudio 5 Real-Time PCR system (Thermo Fisher) cycling conditions were as follows: an initial step (95 °C for 10 min), preceded by 40 cycles of two-step PCR (95 °C for 15 s, 60 °C for 1 min), and then a dissociation step (95 °C for 15 s, 60 °C for 1 min), and a sequential increase to (95 °C for 15 s). Relative mRNA expression was calculated using the ΔΔCt method and glyceraldehyde 3-phosphate dehydrogenase was used as a reference RNA.

### In situ hybridization

In situ hybridization was performed using the RNAscope® Multiplex Fluorescent Reagent kit version 2 (Advanced Cell Diagnostics) according to the manufacturer’s instructions. Images were captured on Nikon 90i upright microscopes (Nikon) with Photometrics CoolSNAP-ES monochrome cameras (Roper). Probes were designed to detect *Tfap2c* (860171, NM_201420.2, target region: 696-1733), *Krt8* (87304-C2, NM_199370.1, target region: 134-1472) and *Prl7b1* (860181-C2, NM_153738.1, target region: 28-900).

### Immunohistochemistry

Frozen rat placentation sites were sectioned at 10 μm, mounted on slides, and fixed in 4% paraformaldehyde. Sections were blocked with 10% goat serum (50062Z, Thermo Fisher) and incubated overnight at 4°C with a primary antibody for vimentin (1:300, sc-6260, Santa Cruz Biotechnology), pan-cytokeratin antibody (1:100, F3418, Sigma-Aldrich), perforin (1:300, TP251, Amsbio), or TFAP2C (1:150, 2320, Cell Signaling Technology). Fluorescence-tagged secondary antibodies, Alexa Fluor 488-conjugated goat anti mouse IgG (A11001, Thermo Fisher), and Alexa Fluor 568-conjugated goat anti-rabbit IgG (A11011, Thermo Fisher) were used to visualize immunostaining.

Sections were counterstained with 4,6-diamidino-2-phenylindole (DAPI, 1:50,000, D1306, Invitrogen). Slides were mounted in Fluoromount-G (0100-01, SouthernBiotech) and imaged on Nikon 90i upright microscopes with Photometrics CoolSNAP-ES monochrome cameras (Roper). Regions of the uterine-placental interface possessing cytokeratin-positive cells were quantified using ImageJ software, as previously described by our laboratory (63, 64).

### Western blotting

Tissue lysates were processed using the radioimmunoprecipitation assay lysis buffer (sc-24948A, Santa Cruz Biotechnology) and protein concentrations determined using the DC Protein Assay Kit (5000112, Bio-Rad). Proteins (80 µg/lane) were separated by sodium dodecyl sulfate polyacrylamide gel electrophoresis. Separated proteins were electrophoretically transferred to polyvinylidene difluoride membranes (10600023, GE Healthcare) for 1 h at 25 V on a semi-dry transfer apparatus (Bio-Rad). Membranes were subsequently blocked with 5% milk for 1 h at room temperature, followed by incubation with antibodies to TFAP2C (1:100, sc-12762, Santa Cruz Biotechnology) and GAPDH (1:5,000, AM4300, Invitrogen) in 5% milk overnight at 4°C. After primary antibody incubation, the membranes were washed in Tris-buffered saline with Tween 20 (TBS-T) three times (10 min/wash) at room temperature. The membranes were then incubated with anti-mouse immunoglobulin G conjugated to horseradish peroxidase (HRP; 1:5000, Cell Signaling Technology) in 5% milk for 1 h at room temperature, washed in TBS-T three times (10 min/wash) at room temperature, immersed in Immobilon Crescendo Western HRP Substrate (WBLUR0500, Sigma-Aldrich), and luminescence detected using Chemi Doc MP Imager (Bio-Rad).

### Identification of TFAP2C and potential TFAP2C gene targets in rat invasive trophoblast cells

Uniform manifold approximation and projection (**UMAP**) plots for *Tfap2c* and *Prl7b1* were generated from rat gd 19.5 uterine-placental interface single cell RNA sequencing (accession No. GSE206086, (38)). Gene expression profiles and open chromatin containing TFAP2C DNA binding motifs identified in rat invasive trophoblast cells (38, 41) were integrated and ascribed function through manual annotation using the UniProt (https://www.uniprot.org) and National Center for Biotechnology (https://www.ncbi.nlm.nih.gov/) databases.

### Statistics

Statistical analyses were performed with GraphPad Prism 10.2.3 software. Statistical comparisons were evaluated using Student’s *t* test 2-tailed or one-way ANOVA with Tukey’s or Dunnett’s post hoc tests as appropriate. Statistical significance was determined as P<0.05.

### Study approval

All protocols using rats were approved by the University of Kansas Medical Center Animal Care and Use Committee, Kansas City, Kansas (approved protocol 22-01-220).

### Data available

All data and materials for this manuscript are included in the Methods and Supporting Data.

## Author Contributions

E.M.D., K. I., and M.J.S. conceived and designed the research; E.M.D. and A.M-I., performed experiments; K.I. provided experimental tools for analysis; E.M.D., A.M-I., R.L.S, and M.J.S. analyzed the data and interpreted results of experiments; E.M.D. and M.J.S. prepared the manuscript. All authors read, contributed to editing, and approved the final version of manuscript.

## Acknowledgments

We thank Brandi Miller, Stacy Oxley, and Leslie Tracy for administrative assistance. The work was supported by the Lalor Foundation (E.M.D., A.M.-I.), Kansas Idea Network of Biomedical Research Excellence, P20 GM103418 (E.M.D, A.M.-I.), NIH grants: HD115834 (A.M.-I.), HD104495 (R.L.S.), HD104071 (K.I.), HD020676 (M.J.S.), HD105734 (M.J.S.), HD112559 (M.J.S.), and the Sosland Foundation (M.J.S.).

## Supplemental information

Figures S1 to S6 and Table S1 to S7.

Dataset S1. Excel file.

