## Supplemental material for "TFAP2C is a key regulator of intrauterine trophoblast cell invasion and deep hemochorial placentation"

**Supporting Figures and Tables**

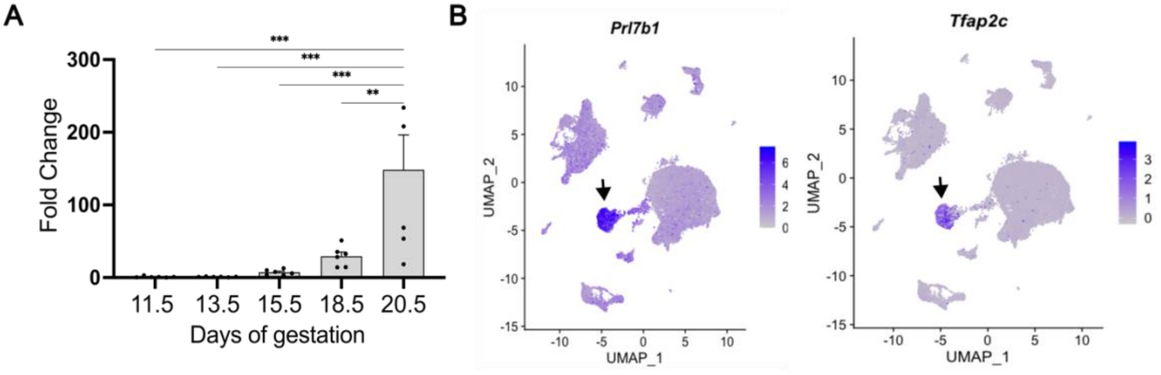


**Supplemental Figure 1. *Tfap2c* transcript expression in the rat uterine-placental interface. A)** RT-qPCR of *Tfap2c* transcript in uterine-placental interface tissues on gestation days 11.5, 13.5, 15.5, 18.5, and 20.5. Data are expressed as mean ± standard error of the mean. Each data point represents a biological replicate from six different pregnancies (n=6). Unpaired *t*-test: **p<0.01, ***p<0.001. **B**) Uniform manifold approximation and projection plot (**UMAP**) for *Prl7b1* and *Tfap2c* transcripts from single cell RNA sequencing of the rat uterine-placental interface at gestation day 19.5 (38). Note the co-localization of *Prl7b1* and *Tfap2c* transcripts in the invasive trophoblast cell cluster (arrows).

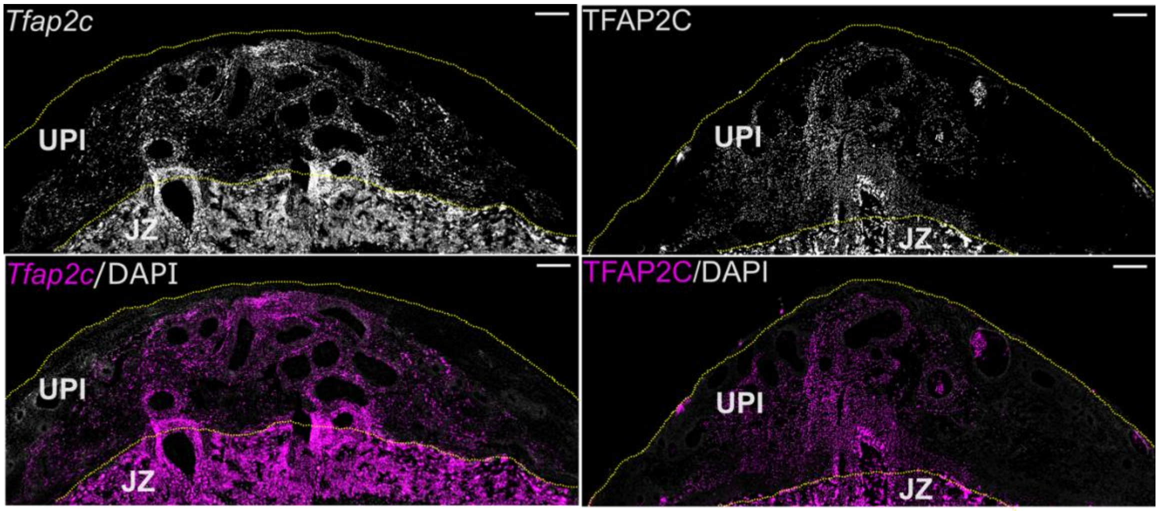


**Supplemental Figure 2. TFAP2C transcript (left panel) and protein (right panel) localization within the gestation day 18.5 uterine-placental interface.** TFAP2C positive cells (magenta) are distributed within the junctional zone (JZ) and uterine-placental interface (UPI). Abbreviation. Scale bar: 500 μm.

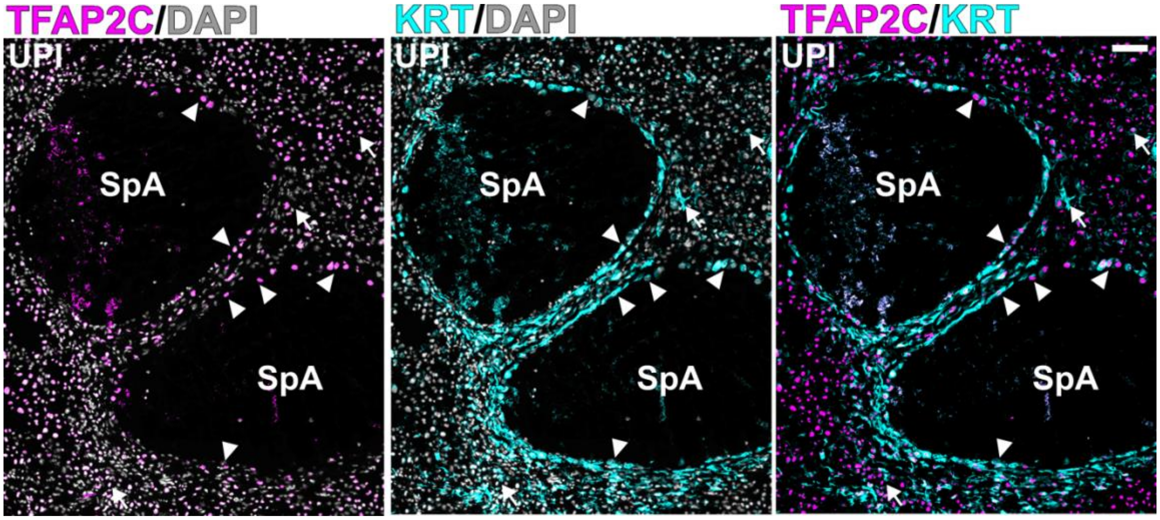


**Supplemental Figure 3. TFAP2C protein localization in the uterine-placental interface.** Immunohistochemistry of TFAP2C (magenta) and cytokeratin (KRT, cyan) within the rat uterine-placental interface at gd 18.5. TFAP2C was localized in nuclei of both endovascular (arrowhead) and interstitial (arrow) invasive trophoblast cells. Abbreviations, UPI, uterine-placental interface; SpA, spiral artery. Scale bar: 100 μm

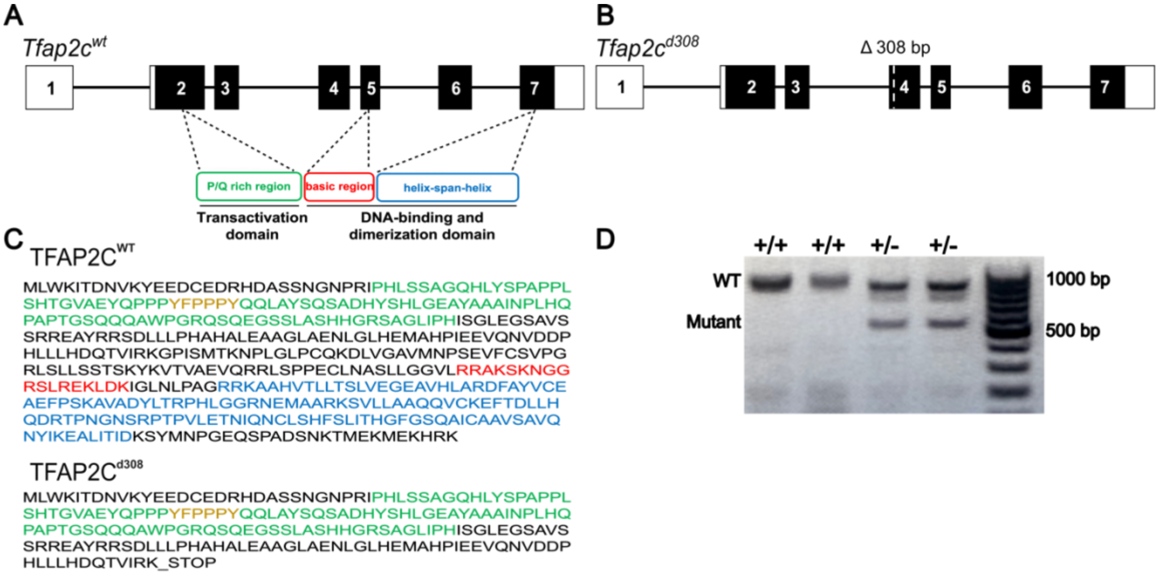


**Supplemental Figure 4. Global disruption of *Tfap2c*. A**) Schematic of *Tfap2c* exon-intron structure and encoded functional domains within the TFAP2C protein. **B**) Schematic of the *Tfap2c* gene possessing a 308 bp deletion (*Tfap2c*$^{d308}$). **C**) Amino acid sequences corresponding to wild type TFAP2C (**TFAP2C$^{WT}$**) and the amino acid sequence for mutant TFAP2C (**TFAP2C$^{d308}$**). **D**) Genotyping of *Tfap2c*$^{WT}$ and heterozygous *Tfap2c*$^{d308}$ conceptuses.

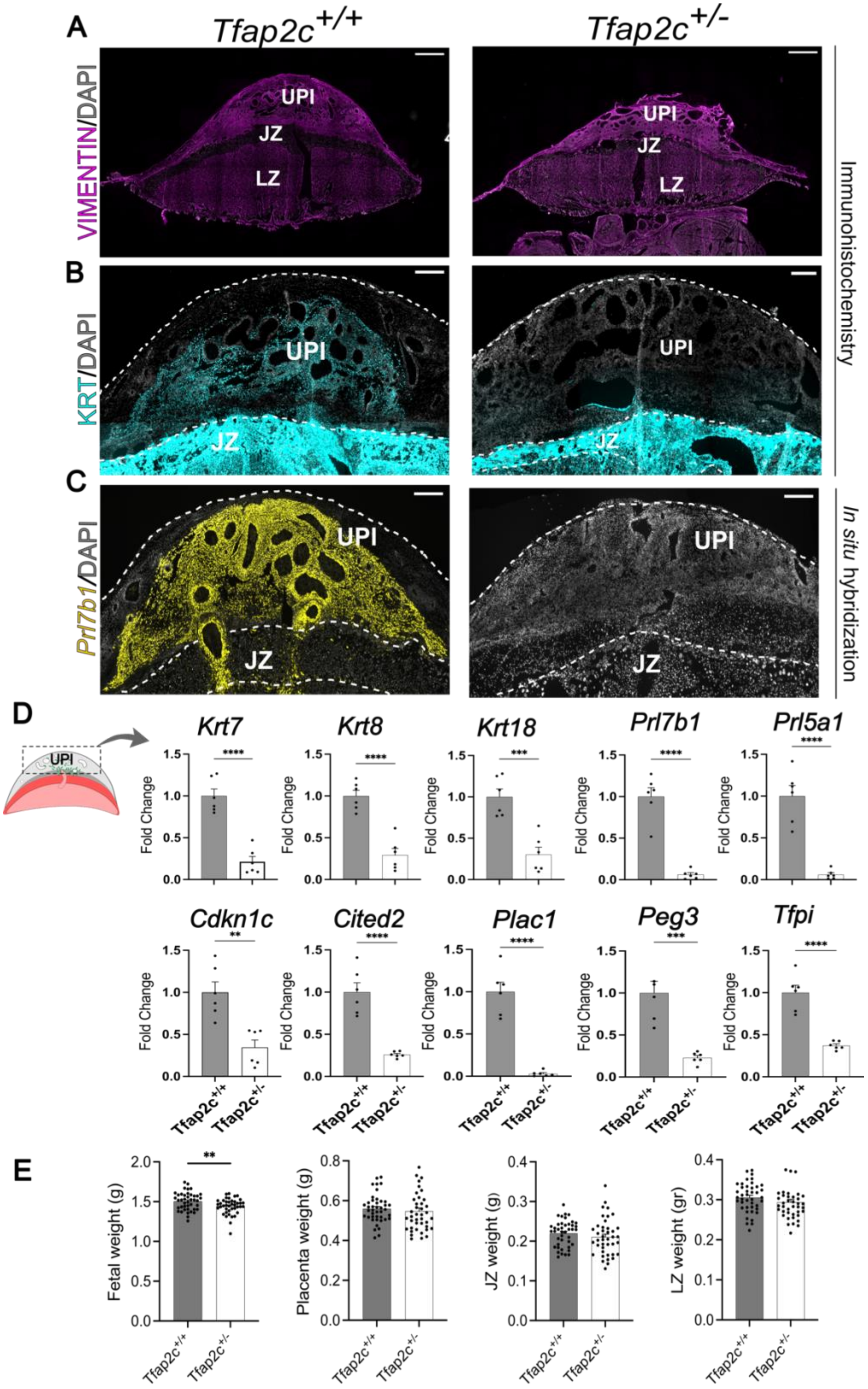


**Supplemental Figure 5. *Tfap2* gene dosage effects on placenta development.** Placentation sites were generated from female *Tfap2c*$^{+/-}$ x male *Tfap2c*$^{+/+}$ breeding. **A**)

Identification of placentation site compartments using vimentin immunostaining on gestation day (**gd**) 18.5 *Tfap2c*$^{+/+}$ and *Tfap2c*$^{+/-}$ placentation sites (magenta). **B**) Immunohistochemical localization of cytokeratin protein in gd 18.5 *Tfap2c*$^{+/+}$ and *Tfap2c*$^{+/-}$ placentation sites (cyan). **C**) Distribution of *Prl7b1* transcripts in gd 18.5 *Tfap2c*$^{+/+}$ and *Tfap2c*$^{+/-}$ placentation sites (yellow). **D)** RT-qPCR of invasive trophoblast cell-specific transcripts in gd 18.5 *Tfap2c*$^{+/+}$ and *Tfap2c*$^{+/-}$ uterine-placental interface tissues. Data are expressed as mean ± standard error of the mean (**SEM**). Each data point represents a biological replicate from six different pregnancies (n=6). **E**) Fetal, placenta, junctional zone and labyrinth zone weights from gd 18.5 *Tfap2c*$^{+/+}$ and *Tfap2c*$^{+/-}$ conceptuses. Data are expressed as mean ± SEM. Each data point represents a biological replicate from six different pregnancies (*Tfap2*+$^{+/+}$, n=41; *Tfap2c* $^{+/-}$, n=42). Unpaired *t*-test: **p<0.01, ***p<0.001, p<0.0001. Abbreviations: UPI, uterine-placental interface; JZ, junctional zone; LZ, labyrinth zone; SpA, spiral artery. Scale bars: 500 μm.

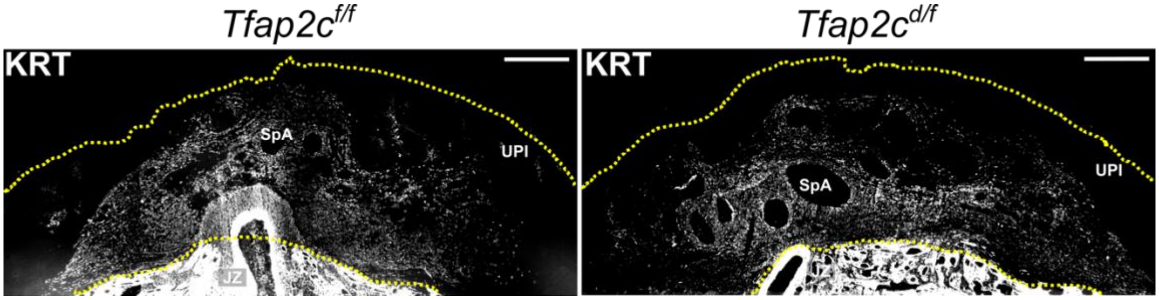


**Supplemental Figure 6. Invasive trophoblast cell distribution within the uterine-placental interface of the *Tfap2c*$^{f/f}$ and *Tfap2c*$^{d/f}$ placentation sites.** Invasive trophoblast cells were localized in the uterine-placental interface using immunocytochemistry for cytokeratin (white). Abbreviations, SpA, spiral artery; UPI, uterine-placental interface. Scale bar: 500 $\mu$m.

**Supplemental Table 1. Genotyping results from female *Tfap2c*$^{+/-}$ x male *Tfap2c*$^{+/-}$ matings.**

| Samples | *Tfap2c* genotype | | |
|---|---|---|---|
| Gestation day: | +/+ (%) | +/- (%) | -/- (%) |
| 8.5 | 3 (14) | 13 (59) | 6 (27) |
| 9.5 | 7 (22) | 25 (78) | 0 (0) |
| 12.5 | 6 (32) | 13 (68) | 0 (0) |
| 15.5 | 23 (41) | 33 (59) | 0 (0) |
| 18.5 | 20 (34) | 38 (66) | 0 (0) |
| Postnatal day 1 | 23 (48) | 25 (52)* | 0 (0) |

*3 *Tfap2c*$^{+/-}$ pups found dead.

**Supplemental Table 2. Genotyping results from female *Tfap2c*$^{+/-}$ x male *Tfap2c*$^{+/+}$ matings.**

| Samples | *Tfap2c* genotype | |
|---|---|---|
| Gestation day: | +/+ (%) | +/- (%) |
| 15.5 | 38 (46) | 44 (54) |
| 18.5 | 41 (51) | 40 (49) |
| Postnatal day 1 | 24 (53) | 21 (47) |

**Supplemental Table 3. Genotyping results from female *Tfap2c*$^{+/+}$ x male *Tfap2c*$^{+/-}$ matings.**

| Samples | *Tfap2c* genotype | |
|---|---|---|
| Gestation day: | +/+ (%) | +/- (%) |
| 15.5 | 33 (60) | 22 (40) |
| 18.5 | 34 (62) | 28 (45) |
| Postnatal day 1 | 22 (61) | 14 (39) |

**Supplemental Table 4. Guide RNAs used for genome editing.**

| gRNA | DNA sequence |
| --- | --- |
| *Tfap2c (Exon4)* | GGTTCATGACCGCTCCAACTGTTTTAGAGCTATGCT |
| *Tfap2c (5' of Exon 4)* | TTAAGTTATATACACTGACAGTTTTAGAGCTATGCT |
| *Tfap2c (3' of Exon 4)* | TTAAGGCCGCAGCTGCATGGGTTTTAGAGCTATGCT |

**Supplemental Table 5. Template sequences used for genome editing.**

| |
|---|
| *Tfap2c 5' loxP template (5' of Exon 4)* |
| CTTAATGAGGGGTTTGTAGGAGGGAGGAGATGGGGGGAATCGGATTACAGT<br>GAGTGGGTGGCTCTTGCTTAAGTTATATACACTGATAACTTCGTATAGCATAC<br>ATTATACGAAGTTATACACGGTTACCGGATATGGAATCGCCCTAAATTACAG<br>GTAGTTATGGTCTTGAGATTGTGAACTGTATTAGTAAATTGGGA |
| *Tfap2c 3' loxP template (3' of Exon 4)* |
| TTCTTTTTGAAAGTGCCAAAATGCAGAAGGTGCTGAAACAGCATAACAGGTT<br>ATCATTTGGTTGGGATTAAGGCCGCAGCTGCAATAACTTCGTATAGCATACA<br>TTATACGAAGTTATTGGAGGGGAGGGGTGGCTTCGAGTGTTTGGTCCCCCT<br>GGGGGAGTTTACTGCTACAATCCTAAGTGAAGATTTTTCTGCACC |

**Supplemental Table 6. Primer sequences used for genotyping.**

| Primer Name | Forward Primer | Reverse Primer |
|---|---|---|
| *Tfap2c* | AGGGGGAGGATGCCATTTAT | GGGGACCAAACACTCGAAG |
| *5' Floxed Tfap2c* | GGGTTTGTAGGAGGGAGGAG | GGTCTGCTCAGAAACAGTTTTC |
| *3'Floxed Tfap2c* | AAGGTGCTGAAACAGCATAACA | GGCCTCCATTTTTGGATTTC |
| *Prl7b1Cre* | TGCTGGAAGATGGCGATTAG | GGTTCCCAGCAAAGTACCAC |
| *Kdm5d* | TTGGTGAGATGGCTGATTCC | GGTTTCTTAAACCGTCGCC |
| *Kdm5c* | TTTGTACGACTAGGCCCCAC | GGTTTCTTAAACCGTCGCC |

**Supplemental Table 7. Primer sequences used for RT-qPCR.**

| Primer Name | Forward Primer | Reverse Primer |
|---|---|---|
| *Peg3* | AAGTTCACGTCCACTCCGTC | CGTCTGGTCTTGTTCGTGGA |
| *Plac1* | CCGTCTCTCCAGATGTCGTT | GAGCCCTTGGAAGCATAGTG |
| *Cited2* | TCTTGGCTGCATGAACTTTG | GGGAGACAGCCAACTTGAAA |
| *Cdkn1c* | CAAACGTCTGCGATGAGTTAGT | AGCCGAAGCCCAGAGTTC |
| *Tfpi* | GCCCGAGGAAGACGATGATA | TCCGCCTTCATTGCACAG |
| *Prl5a1* | TCCACACCAGACATTCCAGA | TTTCCAGGAAGCCAACATTC |
| *Prl7b1* | CCGTCATACTGTCTCAGCACATC | AGCTGTTGAGACCATTGACAACAA |
| *Gapdh* | GACATGCCGCCTGGAGAAAC | AGCCCAGGATGCCCTTTAGT |
| *Krt7* | CGGAATGGGACCTGTGAA | GTAGATGTGGTCTTGATGGAATAGG |
| *Krt8* | TGGGCCAGGAGAAGCTGAA | CACATCCTTCTTGATGAGGACAAA |
| *Krt18* | CTGGAAACCGAGAACAGGAGA | CGGGCATTGTCCACAGAA |
| *Prf1* | GGCACTCAAGAACCTTCC | CTCAAGCAGTCTCCTACC |
| *Klrb1c* | GTCTCCAGGGCATAAGCAAG | AAGAAGGATCAGCGTAGCACA |
| *Klrb1a* | CGTTCACACAGGTTGGCTTT | TGGCTCCACTGATGGTTTTT |
| *Klrb1* | GGTCTCGCTGACTGTTCTTTG | TTGTTGTCCTTTTCCCTTTG |
| *Ncr1* | ATGGGAACATCCAAGCAGAG | ACAGGCTCACTGGGAAAAGA |
